# A novel dataset of 2,362 equine fecal microbiomes from eight veterinary teaching hospital on three continents reveals dominant effects of geography, breed, and disease

**DOI:** 10.1101/2024.10.21.619412

**Authors:** Zachary L. McAdams, Emma J. Campbell, Rebecca A. Dorfmeyer, Giedre Turner, Samantha Shaffer, Tamara Ford, Jenna Lawson, Jackson Terry, Murugesan Raju, Lyndon Coghill, Lucia Cresci, Kara Lascola, Tiffany Pridgen, Anthony Blikslager, Emily Barrell, Heidi Banse, Linda Paul, Alexandra Gillen, Sascha Nott, Marie VandeCandelaere, Gaby van Galen, Kile S. Townsend, Lynn M. Martin, Philip J. Johnson, Aaron C. Ericsson

## Abstract

Horses and other equids are reliant on the gut microbiome for health, and studies have reported associations between certain clinical conditions and features of the fecal microbiome. However, research to date on the equine fecal microbiome has often relied on small sample sizes collected from single and relatively localized geographic regions. Previous work largely employs single timepoint analyses, or horses selected based on limited health criteria. To address these issues and expand our understanding of the core microbiome in health, and the changes associated with adverse outcomes, the Equine Gut Group (EGG) has collected and performed 16S rRNA sequencing on 2,362 fecal samples from 1,190 healthy and affected horses. Here we present the EGG database and demonstrate its utility in characterizing the equine microbiome in health and acute gastrointestinal disease. The EGG 16S rRNA database is a valuable resource to study the equine microbiome and its role in equine health.

## Main Text

With comparatively small stomachs and voluminous hindguts, horses (and other members of the *Equidae* family) tend to eat throughout the day, and continuous throughput of digesta is essential to provide cellulose and other indigestible carbohydrates as substrates for fermentation by hindgut bacteria^1,2^. While the equine gut microbiome represents a rich source of potential diagnostic or prognostic information, most research investigating relationships between the fecal microbiome and clinical features relies on relatively small sample sizes. Owing to their size and the challenges associated with transport, visits to veterinary hospitals are less frequent for horses compared to other domesticated animals, making subject accrual difficult. Moreover, horses typically spend a substantial portion of their life outdoors and geographic and environmental factors likely influence the macromolecular content of dietary substrates, and the microbiome composition. While the effect of geography on the equine fecal microbiome among healthy horses within the context of the same continent appears minimal^3^, recent work has revealed significant differences between healthy domestic horses in Finland, Spain, and Argentina^4^ or in different states within the United States (US)^5^. However, like much equine research, these findings are based on relatively small sample sizes from each location.

One of the most common complaints related to the digestive health of equids is colic, a syndrome of abnormal behavior and other consequences of pain resulting from gastrointestinal obstruction. While several etiologies (e.g., colitis, mechanical obstruction, dysmotility) and risk factors are well-recognized^6^, a substantial portion of cases go undiagnosed and are attributed to nonspecific dysbiosis, or aberrant compositional changes in the gut microbiome^1^. Investigations of the fecal microbiome in horses affected with colic have identified a number of consistent and characteristic changes in the fecal microbiome, including reduced richness^5,7,8^ (i.e., the number of species in a community) and enrichment of certain phyla including *Pseudomonadota*^5,9,10^. While these studies include healthy controls or longitudinal study designs, our knowledge regarding the microbiome of healthy horses, horses affected with gastrointestinal disease, and horses affected with other inflammatory conditions is limited. Other common inflammatory conditions in horses such as laminitis, equine metabolic syndrome, and obesity have also been associated with dysbiosis characterized by reduced microbial richness^11^ and increased relative abundance of *Pseudomonadota*^12,13^. While these findings are not universal^14,15^, this suggests the likely impact of other unrecognized variables. It is unclear how disease status and geographical location, along with factors such as sex, age, and breed interact to influence the equine fecal microbiome.

To address these knowledge gaps and provide a data resource for researchers investigating the equine microbiome, we established the Equine Gut Group (EGG) comprised of researchers at eight veterinary teaching hospitals on three continents, including the University of Missouri (MU), Auburn University, Louisiana State University, North Carolina State University, and University of Minnesota in the US; the University of Liverpool in the United Kingdom (UK); and the University of Queensland and University of Sydney in Australia (AUS). At each institution, fecal samples were collected from subjects admitted to the hospital for any reason, including healthy horses admitted for reasons unrelated to clinical disease. Subject demographics (age, sex, and breed) and detailed medical records accompany these publicly available data representing nearly 2,400 separate fecal samples from over 1,100 individual equine hosts. Here we describe the EGG resource and demonstrate its utility by presenting two vignettes that characterize the healthy adult equine microbiome and compare them to the adult equine colic microbiome.

## Results

### Subject demographics

A total of 2,437 fecal samples were collected from 1,241 unique equine hosts and shipped to the MU Metagenomics Center for processing and 16S rRNA sequencing. After removing samples with uninformative metadata and those yielding less than 10,000 sequencing reads after 16S rRNA sequencing, a total of 2,345 samples from 1,177 horses (*Equus caballus*), 10 samples from 8 donkeys (*Equus asinus*), 6 samples from 4 mules (*Equus asinus × Equus caballus*) and one sample from a single hinny (*Equus caballus × Equus asinus*) were included in the EGG dataset. Most samples were collected from equids in the US; primarily from the Midwest and Southeast regions near donating institutions (**Fig. 1a**, **Table 1**). The University of Missouri donated the largest number of samples (1,218 samples from 707 horses), representing over two-thirds of counties in Missouri (**Extended Data Fig. 1**). All samples were collected between 2015 and 2022, and across all 12 months (**Fig. 1b-c**).

**Fig. 1.**
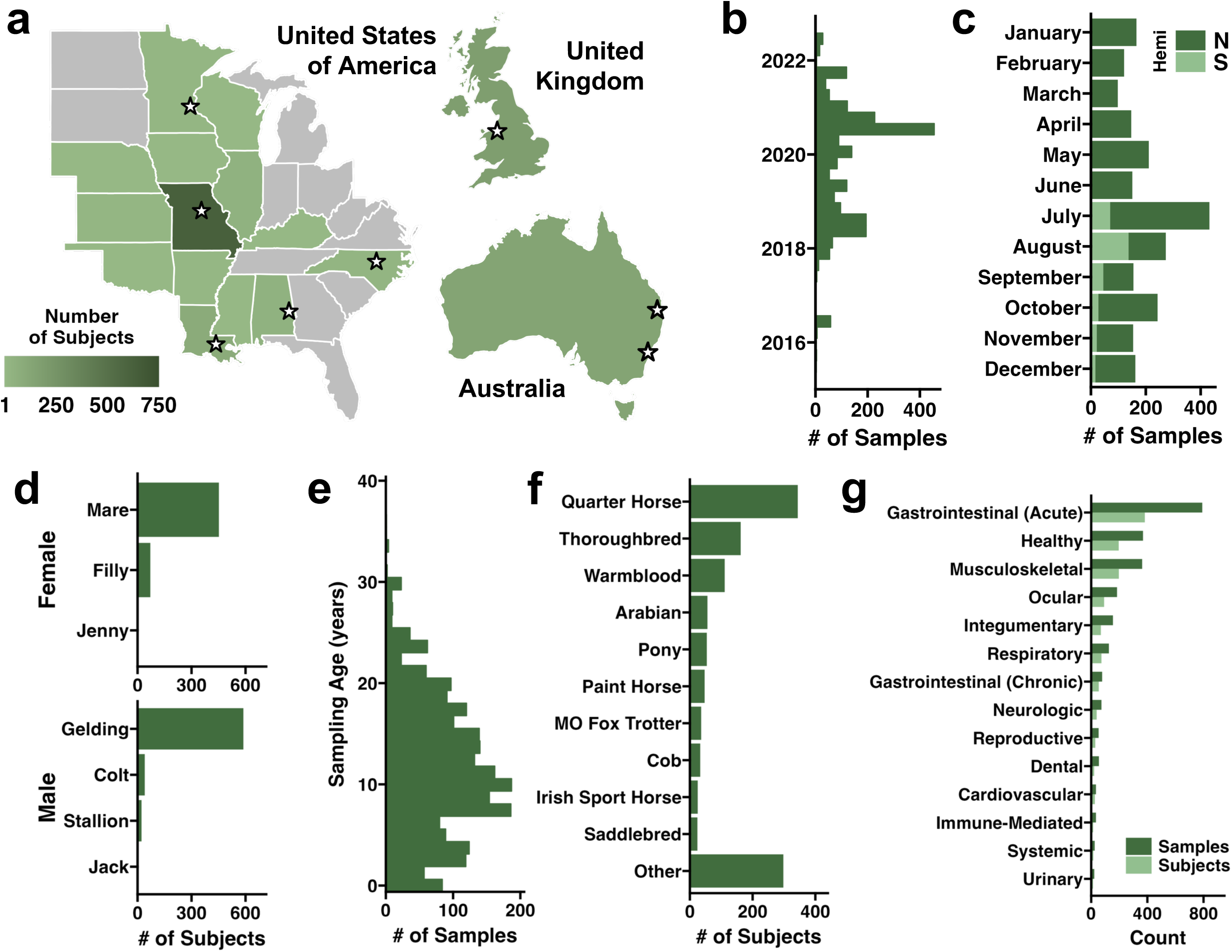
Demographic summary of the EGG database. **a**, Maps representing the number of subjects represented in the EGG resource by state (USA) or country (UK and AUS). Stars indicate the location of donating institutions. **b**, Histogram depicting sample collection date between July of 2015 and July of 2022. **c**, Stacked bar chart depicting the distribution of samples collected during each month in either the northern (USA and UK) or southern (AUS) hemispheres (Hemi). **d**, Bar chart depicting number of male and female subjects separated by breeding status. **e**, Histogram depicting the distribution of patient ages at the time of sampling. **f**, Bar chart depicting the ten most abundant breeds sampled in the dataset. Missouri Fox Trotter shortened to ‘MO Fox Trotter’. All other breeds were grouped together as ‘Other’. **g**, Bar chart depicting the number of samples and donating subjects representing each primary diagnosis category.

Regarding sex and breeding status, the majority of samples were collected from geldings (*n* = 588 subjects) and mares (*n* = 452), with fewer fillies, colts, stallions, jacks, and a jenny sampled (n = 71, 40, 22, 2, and 1, respectively). Overall, 54.8% and 44.0% of subjects were male and female, respectively (**Fig. 1d**). Subjects ranged in age from less than 24 hours to 36.0 years old with an average age at collection of 11.8 ± 6.8 years old (**Fig. 1e**). Forty-six unique breeds of horses, mules, and donkeys were sampled with Quarter Horses, Thoroughbreds, and Warmbloods representing 51.8% of the overall patient population (**Fig. 1f**).

Given the broad range of breeds represented in the EGG database, we grouped similar breeds into one of seven general breed categories commonly used in equine medicine based on phenotypical morphology and/or temperament (**Extended Data Fig. 2a**): Quarter Horse (*n* = 344 subjects), Hot-blooded breeds (*n* = 219), Warmblood (*n* = 166), Pony (*n* = 88), Draught (*n* = 59), Cob (*n* = 42), Donkey (including Mules and a Hinny, *n* = 15), and Other (*n* = 257).

Given these subjects were admitted to each donating institution for clinical reasons, boarding, or as companions to other subjects, we reviewed the available medical records for each patient and categorized each hospital visit into one of fourteen general categories denoting the primary presenting complaint, while also incorporating secondary categories discovered upon further clinical assessment (**Fig. 1g, Extended Data Fig. 2b**). Longitudinal samples (≥ 3 samples/horse) were collected from 249 (20.4%) healthy and clinical subjects allowing for longitudinal comparisons in health and multiple disease conditions (**Extended Data Fig. 2c**). Reported treatments given to each patient were also recorded individually by generic name and grouped into general categories (e.g., antibiotics, analgesics, etc.; **Extended Data Fig. 2d**). Fatal or owner-requested euthanasia outcomes were reported in 131 (11.0%) subjects. A complete metadata file with the National Center for Biotechnology Information (NCBI) Sequence Read Archive (SRA) accession numbers for all samples included in the EGG resource is provided in **Extended Data Table 1**.

### EGG 16S rRNA Data Summary

Of the 2,437 samples sequenced and processed using DADA2, 2,362 produced high-quality, informative data (> 10,000 reads, **Fig. 2a)**. Total feature counts per sample ranged from 7,762 to 413,897, averaging 92,440 ± 35,227 (**Fig. 2b**). Richness (i.e., Chao1 Index) across all samples ranged from 23 to 2,775, averaging 1,010 ± 360 (**Fig. 2c**). Shannon diversity ranged from 0.82 to 6.7, averaging 5.5 ± 0.87 (**Fig. 2d**). To determine the importance (i.e., increase in mean squared error) of select intrinsic and extrinsic factors influencing alpha diversity, we generated random forest models explaining richness and diversity using the age and sex of the animal, primary clinical diagnosis, geographic location (i.e., country), breed type, and the time of year at which the sample was collected (**Fig. 2e**). Patient age was found to have the highest importance value for both Chao1 and Shannon Indices, however, no significant correlation between age and richness or diversity was observed (**Extended Data Fig. 3a-b**). In addition to richness and alpha diversity, we determined the relative effect size of each of these factors on weighted beta diversity using Bray-Curtis distances. Geographic location (R^2^ = 2.9%) and primary diagnosis category (R^2^ = 2.7%) had the largest effect on beta diversity using the considered metadata factors (**Fig. 2e**). Both alpha and beta diversity were influenced by primary diagnosis category and geographic location (**Extended Data Fig. 3c-h**). Visualizing these differences in beta diversity using principal coordinate analyses (PCoA) revealed clear clustering of samples by both disease category and country of origin supporting the statistical differences found between groups (**Extended Data Fig. 3e, 3h**). Differences in alpha and beta diversity were also observed between donating institutions (**Extended Data Figure 4a-c, Extended Data Table 2**). Samples collected from healthy subjects differed in alpha and beta diversity from multiple primary diagnosis categories including musculoskeletal and ocular conditions (**Extended Data Table 3**). It is important to note that despite the clear separation of groups by disease category and geographic location, the selected factors only explained 6.1% of total variance (Residuals = 92.7%) suggesting other factors may explain more of the variability in beta diversity.

**Fig. 2.**
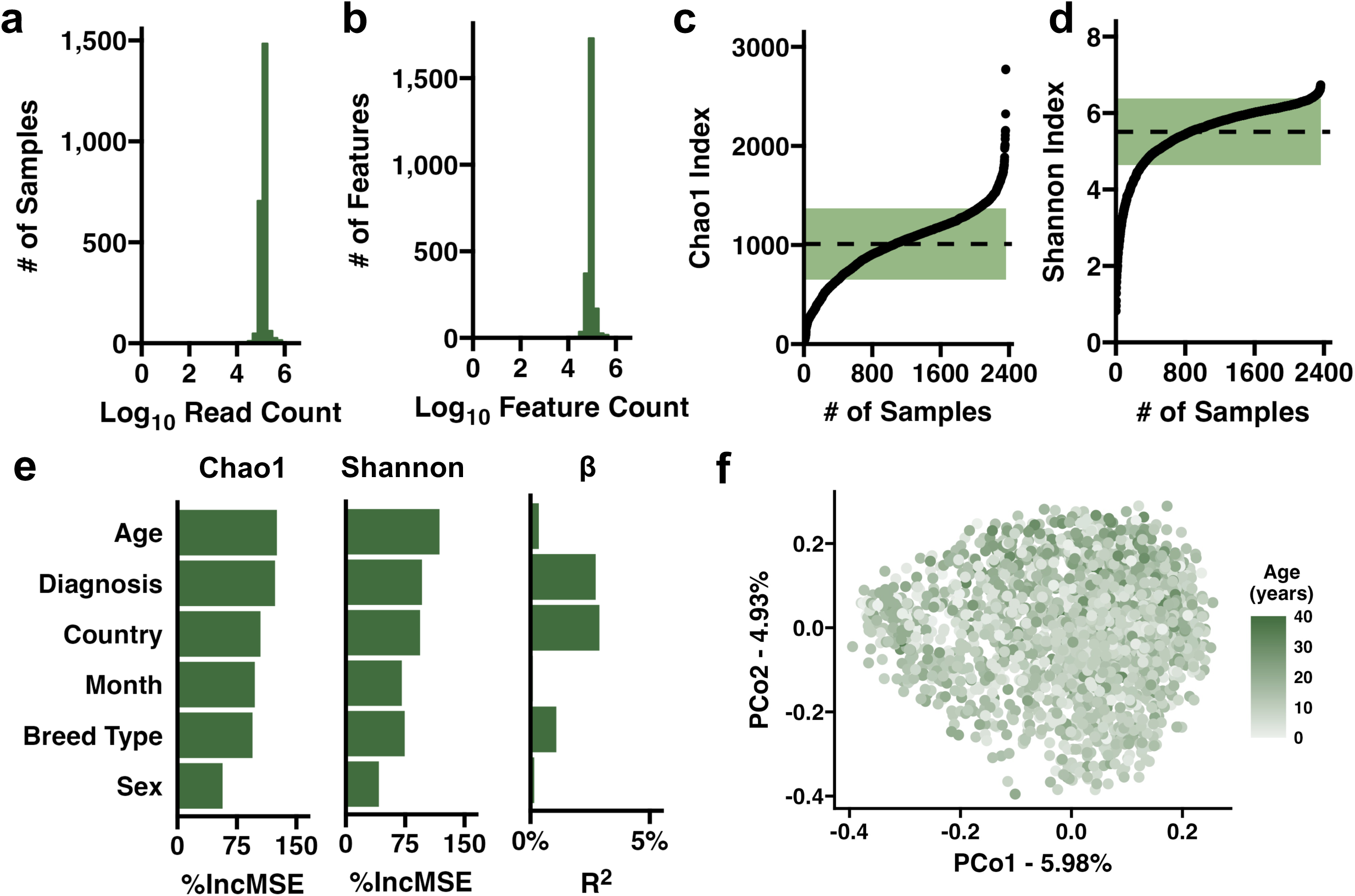
Summary of the EGG 16S rRNA dataset. Histograms depicting the **(a)** number of forward reads and **(b)** unique features observed in each sample. **c**,**d**, Ordered dot plots depicting the observed **(c)** richness and **(d)** diversity in all samples. Dashed line represents the average richness and diversity, respectively. Green bar represents standard deviation. **e**, Bar plots depicting the importance of selected metadata factors influencing overall alpha and beta diversity. Percent increase in mean squared error (%IncMSE) determined by a random forest model identifying factors influencing richness *(left,* 38.98% variation explained) and diversity *(middle,* 40.1% variation explained). Effect size (R^2^) on weighted beta diversity determined using permutational analysis of variance *(right).* **f,** Principal coordinate analysis using weighted distances of all samples depicting patient age at collection. *N* = 2,302 samples.

### Core Microbiome of Healthy Adult Equids

We next characterized the intrinsic and extrinsic factors affecting alpha diversity, beta diversity, and core taxonomic features of the healthy adult (> 4 years old) equine microbiome using a subset of the EGG resource composed of 277 samples collected from 157 subjects (13.7 ± 6.3 years old sampling) of all breed types (**Extended Data Table 4**). Because no differences in richness (*p* = 0.314) or diversity (*p* = 0.690) were observed over time, all healthy samples were included in the following analyses accounting for repeated measures of the same subjects when possible. Of the selected metadata factors (geographic location, breed type, age, sex, time of year sample was collected), random forest models identified geographic location as the most important factor influencing both richness and diversity (**Extended Data Fig. 5a**). Healthy equine subjects from the US exhibited greater richness than healthy adults from both the UK and ASTL (**Extended Data Fig. 5b**). Beta diversity was marginally influenced by geographic location (R^2^ = 4.0%); however, breed type had the largest effect on weighted beta diversity (R^2^ = 7.0%) (**Extended Data Fig. 5a**). Visualizing these differences in beta diversity using PCoA revealed a substantial separation of samples by geographic location along the first principal coordinate (**Extended Data Fig. 5** Samples also clustered by breed type where stocky, cold-blooded draught horses clearly separated from Quarter Horses and other lean, hot-blooded breeds (**Extended Data Fig 5d**).

We then determined the core taxonomic microbiome of the healthy adult equid. Using a prevalence-abundance core microbiome analysis, we identified twenty-one families with a minimum 5% prevalence at 1% relative abundance including twelve families from the phylum *Bacillota*, six from *Bacteroidota*, and one each from *Verrucomicrobiota*, *Spirochaetota*, and *Fibrobacterota* (**Fig. 3a**). WCHB1-41 (phylum *Verrucomicrobiota*), a family positively correlated with microbial diversity in healthy animals and identified as a core member of the equine microbiome^16,17^, exhibited the highest prevalence across relative abundance thresholds in healthy subjects. Other prolific short chain fatty acid (SCFA)producers inducing *Lachnospiraceae* and *Oscillospiraceae* (phylum *Bacillota*) and *Rikenellaceae* (phylum *Bacteroidota*) were highly prevalent in the healthy equine gut being detected at a minimum of 5% relative abundance in more than 83.5% of healthy samples (**Fig. 3a**). Together, these four families made up an average of 42.4% of the healthy equine gut microbiome (**Extended Data Fig. 6**): WCHB1-41 (11.6% ± 6.1%), *Rikenellaceae* (11.1% ± 5.7%), *Lachnospiraceae* (11.1% ± 6.2%), and *Oscillospiraceae* (8.8% ± 4.0).

**Fig. 3.**
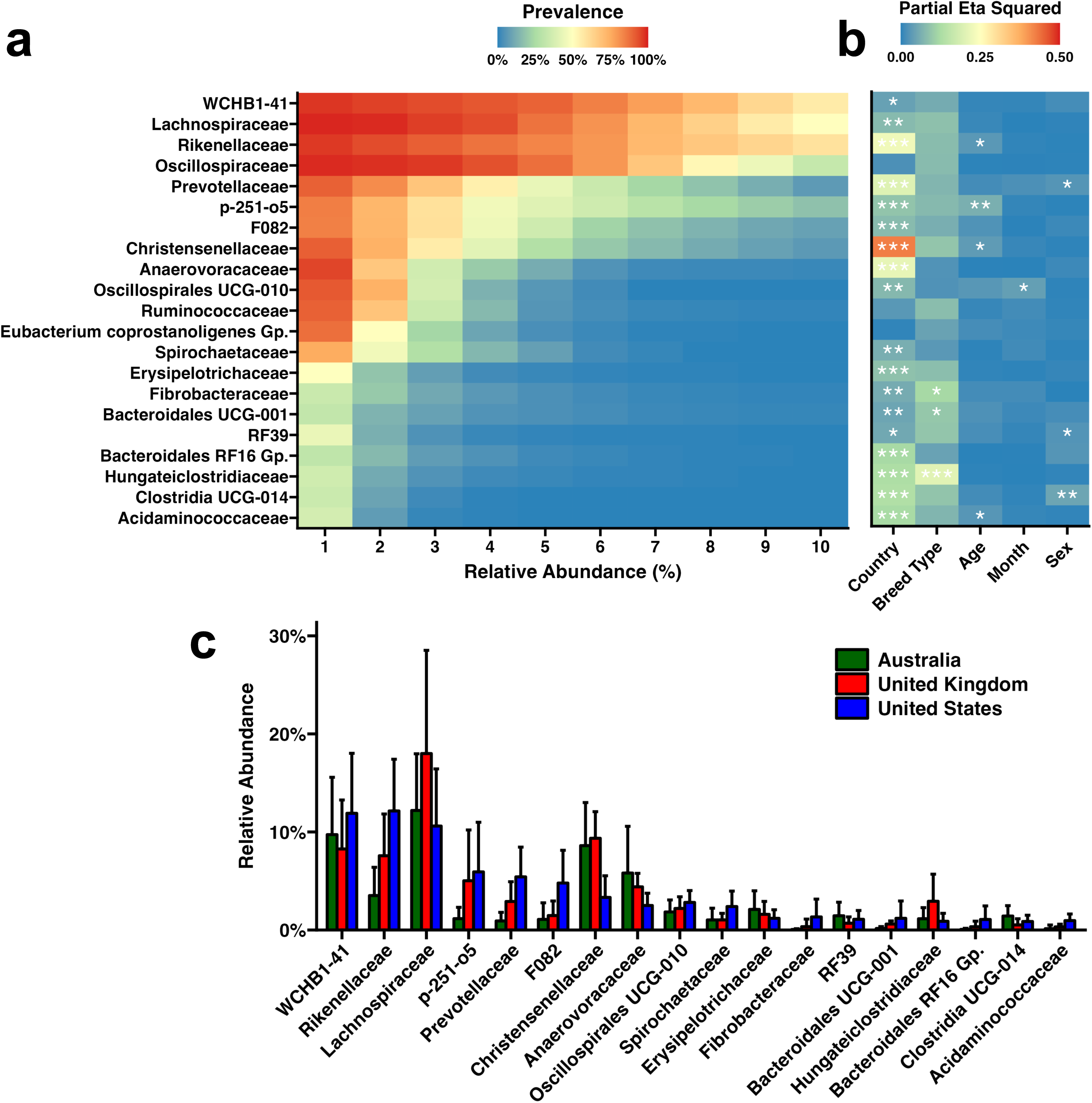
Core microbiome of healthy adult equids. **a**, Heatmap depicting the family-level presence-absence core microbiome of healthy adult equids. **b**, Heatmap depicting significance and effect size of covariates as determined by ANOVA of relative abundance data fitted to linear mixed-effects models for each family. Significance of covariate is indicated within each tile. * *p* < 0.01, ** *p* < 0.01, *** *p* < 0.001. **c**, Bar plots depicting relative abundance of families significantly affected by geographic location. Pairwise Wilcoxon comparisons may be found in Extended Data Table 5.

To further explore host and environmental factors influencing the core microbiome of the healthy equine gut, we fitted relative abundance data for each family to linear mixed-effects models and determined the effect size of age, sex, breed type, geographic location (i.e., country), and the time of year at which the sample was collected. Geographic location significantly affected eighteen families (**Fig. 3b**, **Extended Data Table 5**). When comparing the relative effect size of each factor between families, geographic location had the largest effect on *Christensenellaceae* (η*_p_*^2^ = 0.424) where US subjects exhibited a relative abundance of 3.3% ± 2.2%, nearly three-times lower than subjects from the UK (9.4% ± 2.7%) and AUS (8.6% ± 4.4%) (**Fig. 3c**). Significant effects of breed type, age, month of collection, and sex were observed in eleven of the core families collectively suggesting a multifactorial contribution of intrinsic and extrinsic factors shaping the core taxonomic microbiome of the healthy equine gut (**Fig. 3a-b**).

### Microbial Markers of Acute Gastrointestinal Disease in Equids

We next compared healthy adult subjects to those diagnosed with acute gastrointestinal (GI) disease (i.e., colic).Single timepoint and longitudinal analyses were used to identify microbial markers of acute GI disease using a subset of the EGG resource including 259 samples from 157 healthy equids and 602 samples from 320 colic subjects (including colic as secondary diagnoses; **Extended Data Fig. 2b**). Colic cases were observed in both sexes, at each geographic location, in all breed types, and across a broad range of ages (**Extended Data Fig. 7a-d**). Beginning with a single time point analysis, we assessed differences in community dynamics between healthy and acute GI disease cases at patient intake (Day 1). Decreased richness (*p* = 0.028) and diversity (*p* = 0.003) were observed in acute GI disease subjects compared to healthy animals (**Extended Data Fig. 7e-f**). Beta diversity also differed between disease cases at intake (F = 3.61, *p* < 0.001), however, no clear separation of samples collected from healthy or colic subjects was overserved along the first two principal coordinates (**Extended Data Fig. 7g**).

We then asked whether these differences persisted over the course of one week in the hospital using only subjects from which longitudinal samples (≥ 3 samples from each subjects) were collected (Healthy: 98 samples from 22 subjects; Colic: 270 samples from 70 subjects). Subjects with acute GI disease exhibited increasingly lower richness and diversity over the course of one week (**Fig. 4a, Extended Data Fig. 7h**). PCoA of longitudinal beta diversity suggested marked shifts in microbial composition over time because samples collected from subjects with acute GI disease later in the week separated away from intake samples along the first principal coordinate (**Fig. 4b**). Longitudinal samples collected from healthy subjects remained relatively clustered regardless of time spent in the hospital. This observation was substantiated with a volatility assessment comparing within patient community dissimilarity of longitudinal samples to the intake samples (Day 1). While both healthy and acute GI disease subjects showed significantly increasing dissimilarity from intake samples over time, acute GI disease subjects exhibited double the rate of increasing dissimilarity over time (Slopes: Healthy = 0.032/day, Acute GI = 0.064/day; **Fig. 4c**). Collectively, these single timepoint and longitudinal community-level analyses of the equine microbiome during acute GI disease suggest a microbial dysbiosis that increases with time.

**Fig. 4.**
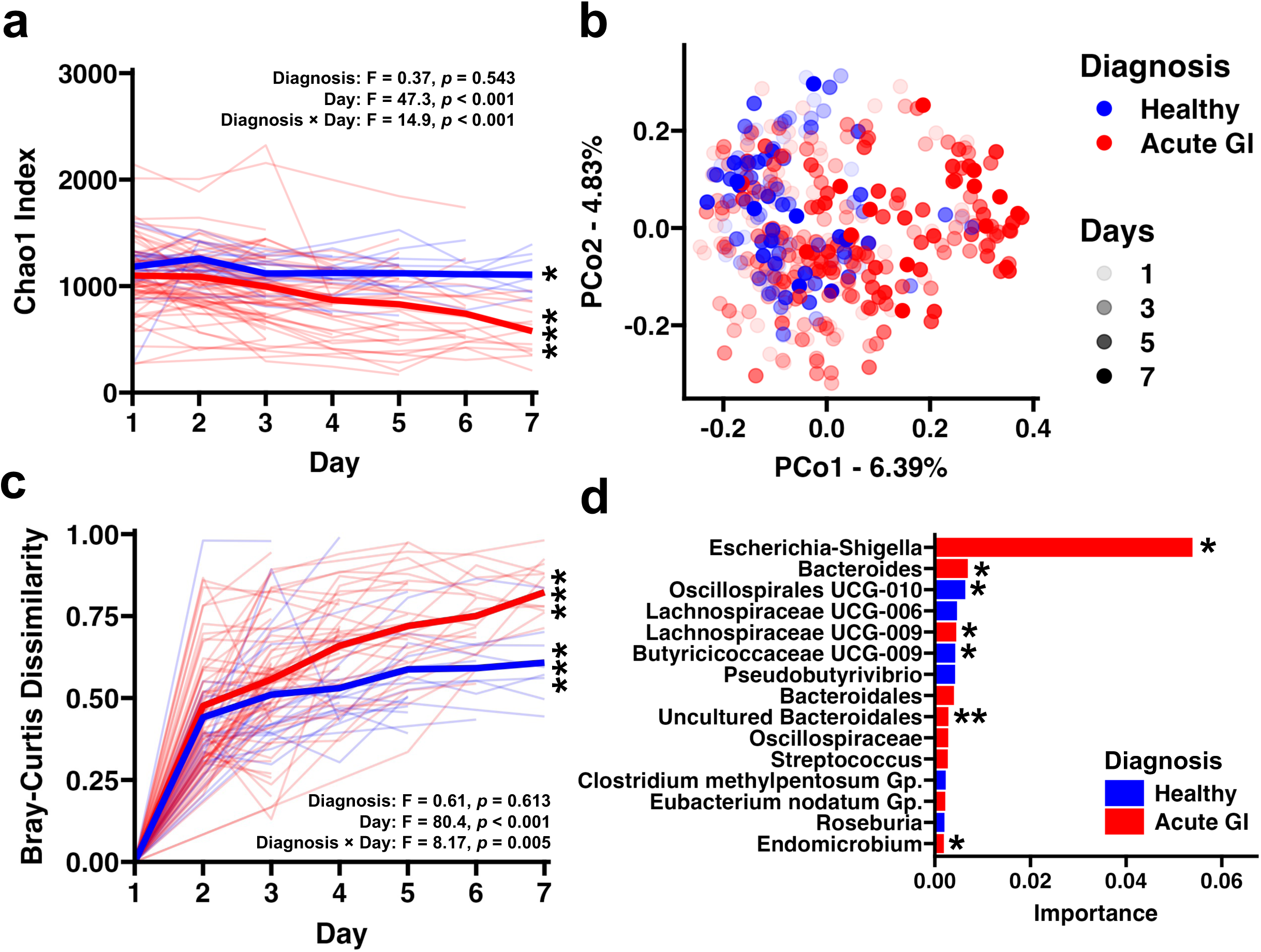
Acute GI disease is characterized by a microbial dysbiosis exacerbated by time. **a**, Line plot showing longitudinal changes in sample richness (Chao1 Index). Thin lines depict individual subjects and thick lines represent group means. Inset text describes two-way ANOVA results of longitudinal data fitted to a LMEM. \**p* < 0.05, \*\*\**p* < 0.001. One-way ANOVA within group. **b**, PCoA using weighted beta diversity showing shifts in microbial composition over time. Opacity of each point represents day of collection. Diagnosis: F = 5.8, *p* < 0.001; Day: F = 3.8, *p* < 0.001; Diagnosis × Day: F = 2.1, *p* < 0.001 **c**, Volatility assessment of microbial composition comparing weighted dissimilarity of each sample to the sample collected at intake from the same patient. Thin lines depict individual subjects and thick lines represent group means. Inset text describes two-way ANOVA results of longitudinal data (Days 2-7) fitted to a LMEM. \*\*\**p* < 0.001. One-way ANOVA within group. **d**, Bar plot showing top 15 taxonomic features by importance in predicting disease status. Bar color depicts which diagnosis category in which the genus was enriched. \**p* < 0.05, \*\**p* < 0.01. *mikropml* permutation testing.

Diagnosing colic in equids is challenging given its diverse and often non-specific etiology. Behaviors and clinical signs, such as looking at the flank or standing with a stretched-out posture, might be associated with extra-intestinal conditions like renal or urinary diseases. Therefore, leveraging microbiome data to predict acute GI disease would improve equine diagnostic testing capabilities and more rapidly target appropriate therapeutic intervention. Using the EGG resource, we generated a random forest model to predict health or acute GI disease with genus-level relative abundance microbiome data using the *mikropml* library^18^. To simulate its practical application in the clinic, we trained and tested the model using microbiome data collected at intake (i.e., Day 1). After pre-processing, relative abundance data from 283 genera were used to create the model. Overall, the model achieved an area under the curve (AUC) of 0.770 (**Extended Data Fig. 8**). While the specificity of this model was low (0.452), sensitivity was high (0.938) suggesting that, in combination with clinical assessments, this model may be used to rule out acute GI disease. When comparing the importance of individual taxonomic features, *Escherichia-Shigella* was found to be the most important predictor of acute GI disease (**Fig. 4d**). SCFA-producers including *Oscillospirales* UCG-010 and multiple *Lachnospiraceae* genera were also important predictors of health or disease (**Extended Data Fig. 7i, Extended Data Table 6**). Collectively, acute GI disease (i.e., colic) is characterized by a robust microbial dysbiosis at the community level. Specifically, reduced richness and substantial shifts in community composition including increased opportunistic pathogens and decreased SCFA-producers are characteristic of colic.

After further review of medical records for subjects with acute GI disease, we identified six subgroup characterizations describing colic etiology; three subgroups of inflammatory conditions and three subgroups of non-inflammatory conditions. Inflammatory etiologies included Mechanical (i.e., intestinal strangulation, entrapment; *n* = 49 subjects), Infectious (i.e., Coronavirus, Salmonellosis, *Neorickettsia risticii*, *Streptococcus equi* ss *equi*, *Clostridioides difficile*, *Rhodococcus equi*, *n* = 19), and Other (i.e., undiffierentiated enteritis, typhlitis, colitis, peritonitis, and inflammatory bowel disease; *n* = 49). Non-inflammatory etiologies included Mechanical (i.e., impaction, enterolith, *n* = 80), Other (i.e., spasmodic colic, gastric glandular disease, flatulent colic; *n* = 33), and Idiopathic (*n* = 97). (**Extended Data Fig. 9a**). Compared to healthy subjects, inflammatory infectious (*p* < 0.001) and non-inflammatory other (*p* < 0.001) but not non-inflammatory mechanical (*p* = 0.440) exhibited lower richness at intake (**Extended Data Fig. 9b, Extended Data Table 7**). Non-inflammatory mechanical also exhibited reduced richness (*p* = 0.035) compared to health subjects at intake. Differences in weighted beta diversity were also observed between healthy and all inflammatory and non-inflammatory subgroups, however, no clear separation of acute GI disease subgroups along the first two principal coordinates was observed (**Extended Data Fig. 9c**, **Extended Data Table 7**).

Given that acute GI disease was characterized by decreased richness and diversity and robust volatility in microbial composition over time (**Figure 4a**, **Figure 4c, Extended Data Fig. 6h**), we next determined if these observations were consistent across acute GI disease subgroups. Healthy and non-inflammatory mechanical, non-inflammatory idiopathic and inflammatory (combining mechanical, infectious, and other) cases were included in this analysis due to sample size limitations of longitudinal data. Given the limited sample size of non-inflammatory idiopathic cases, longitudinal comparisons of acute GI disease subgroups were limited to five days. Richness and diversity significantly decreased over time in inflammatory and non-inflammatory mechanical and idiopathic colic cases (**Extended Data Fig. 9d-e**). While not significant, richness marginally decreased in healthy subjects over time (*p* = 0.090). Lastly, we compared differences in longitudinal beta diversity (**Extended Data Fig. 9f-k**). Significant differences in beta diversity between healthy and inflammatory (F = 5.63, *p* < 0.001), non-inflammatory mechanical (F = 4.62, *p* < 0.001), and non-inflammatory idiopathic (F = 3.55, *p* < 0.001) colic subjects were observed. Visualizing these differences suggested time-dependent shifts in composition in each colic subgroup (**Extended Data Fig. 9f-h**). No differences in volatility were observed between healthy subjects and each colic subgroup yet significant increases in dissimilarity over time were observed in every group (**Extended Data Fig. 9i-k**). Collectively, inflammatory colic cases exhibit a more severe microbial dysbiosis than colic of non-inflammatory mechanical and idiopathic origin.

## Discussion

This catalog of 2,362 informative 16S rRNA sequencing data is a valuable tool for single timepoint and longitudinal retrospective equine microbiota investigations in health and disease. We demonstrated its utility by presenting two unique use cases for the EGG resource. We characterized and defined the core taxonomic microbiome of the healthy adult equine gut and the effect of select intrinsic and extrinsic factors on those features. We then compared the microbiome of healthy equine subjects to that of subjects with acute GI disease identifying both community-level and taxonomic markers of the microbial dysbiosis associated with colic. These examples demonstrate novel applications of the EGG resource and provide evidence for future prospective studies into the restoration and maintenance of equine health via the gut microbiome. Differences in microbial diversity between healthy animals and other disease states were also identified highlighting the broad range of conditions in which the EGG resource may be applied to identify microbial markers of disease.

Core microbiome analyses of the equine gut have previously been limited in sample size and patient diversity^19,20^. While few have incorporated multiple breeds of horses or even multiple species within the *Equidae* family^16^, the effects of host and environmental factors shaping the core microbiome are often underappreciated. Here we identified twenty-one families in the core microbiome of the healthy equid with family-level resolution in a substantially larger and more diverse population of healthy equine subjects. Many of these taxa including WCHB1-41, *Lachnospiraceae,* and *Ruminococcaceae* have previously been identified as members of the core equine microbiome suggesting essential roles for these SCFA producers in the healthy equine gut^16^. In addition to identifying families making up the core microbiome of the healthy equine microbiome, we identified host and environmental factors that influence the relative abundance of these features. Country of origin affected eighteen families suggesting biogeographical factors (e.g., climate) may have robust effects on the equine gut. Having curated the sample collection date and specific geographic location of many subjects in the EGG resource, we have presented precise spatiotemporal coordinates allowing for future retrospective investigations associating meteorological data to microbial diversity in the equine gut.

Having collected fecal samples from subjects with one or more clinical diagnoses in thirteen distinct disease categories, the EGG resource may be used to identify microbial markers across many equine clinical conditions. Here, we identified community-level and taxonomic makers of acute GI disease, or colic. The continuous throughput of fermentable fibers through the equine hindgut supplies the microbiome, and in turn, the host, with much of its energy requirements. Thus, dysmotility of nutrients through the hindgut contributes to both colic pathogenesis and microbial dysbiosis. Our data point to substantial reductions in community richness and diversity and shifts in microbial composition that are exacerbated over time in acute GI disease. We found that these patterns of microbial dysbiosis were consistent across inflammatory and non-inflammatory colic cases suggesting that while multiple mechanisms may contribute to pathogenesis of colic, the resulting gut dysmotility is likely associated with similar patterns of microbial dysbiosis. Measuring these microbiome-specific markers of colic in feces collected from horses in the clinic, however, may be challenging given the time and costs required to generate 16S rRNA sequencing data. Thus, the development of targeted assays measuring the abundance of key taxonomic markers of health or acute GI disease presented here (e.g., *Escherichia-Shigella*, *Lachnospiraceae*) would tremendously benefit veterinary diagnosticians working with equids.

While highly valuable, the EGG resource is not without its limitations. 16S rRNA targeted amplicon sequencing is a financially and computationally advantageous method for characterizing microbial communities of large populations, however, it does not capture the functional capacity (i.e., metagenome) of these microbiomes. Microbes often exhibit functional redundancy meaning taxonomically distinct microbes can fill the same metabolic niche^21^, thus, 16S rRNA sequencing may identify high taxonomic diversity while metagenomic, or even metabolic, diversity is more conserved. Future efforts to characterize the core metagenome (and metabolome) of the equine gut in health and disease may optimize the development of microbial medicines for equine applications. An additional constraint of the EGG resource is the limited resolution of diet-related metadata. Owner-reported diet data were generally restricted to general variations of pasture, grain, hay, and/or nursing. Specific grass or grain content, amounts consumed, and macronutrient contents of these diets were frequently not reported, thus, we were unable to assess these factors despite the known diet-mediated effects on the equine microbiome^4,22^. Lastly, for many subjects in the EGG database, treatment data (i.e., antibiotics, fluids, surgery) was provided, however, precise information as to dosage or timing (including treatments administered by referring veterinarians) were not available, thus, while recognizing the potential effects of these variables on the microbiome, we could not currently incorporate these into our analysis.

The EGG resource is a highly valuable collection of equine fecal microbiome data for the retrospective analysis of the equine gut microbiome in health and disease. It was created to address the major limitations facing the equine microbiome field: insufficient sample sizes and lack of subject diversity. These data represent a diverse equine population from a wide geographic area, integrated with rich metadata, and is intended to provide the veterinary and research communities a useful resource for preliminary *in silico* investigations, and a publicly available reference point for the equine microbiome. As samples are added and the EGG continues to grow, efforts are also underway to develop a NoSQL database and web portal, facilitating access and enhancing utility of the data. With the EGG database in its current form, it provides a tremendous research and clinical resource to investigate equine GI disease and other conditions, improve clinical diagnostics capabilities, and ultimately advance equine health.

## Data availability

All 16S rRNA amplicon sequencing data and associated metadata supporting the present study are available at the National Center for Biotechnology Information (NCBI) Sequence Read Archive (SRA) under BioProject PRJNA1032075. All code is available at https://github.com/ericsson-lab/EquineGutGroup.

## Acknowledgements

We would like to acknowledge the staff and veterinary students at all participating veterinary teaching hospitals.

---

**Extended Data Fig. 1.**
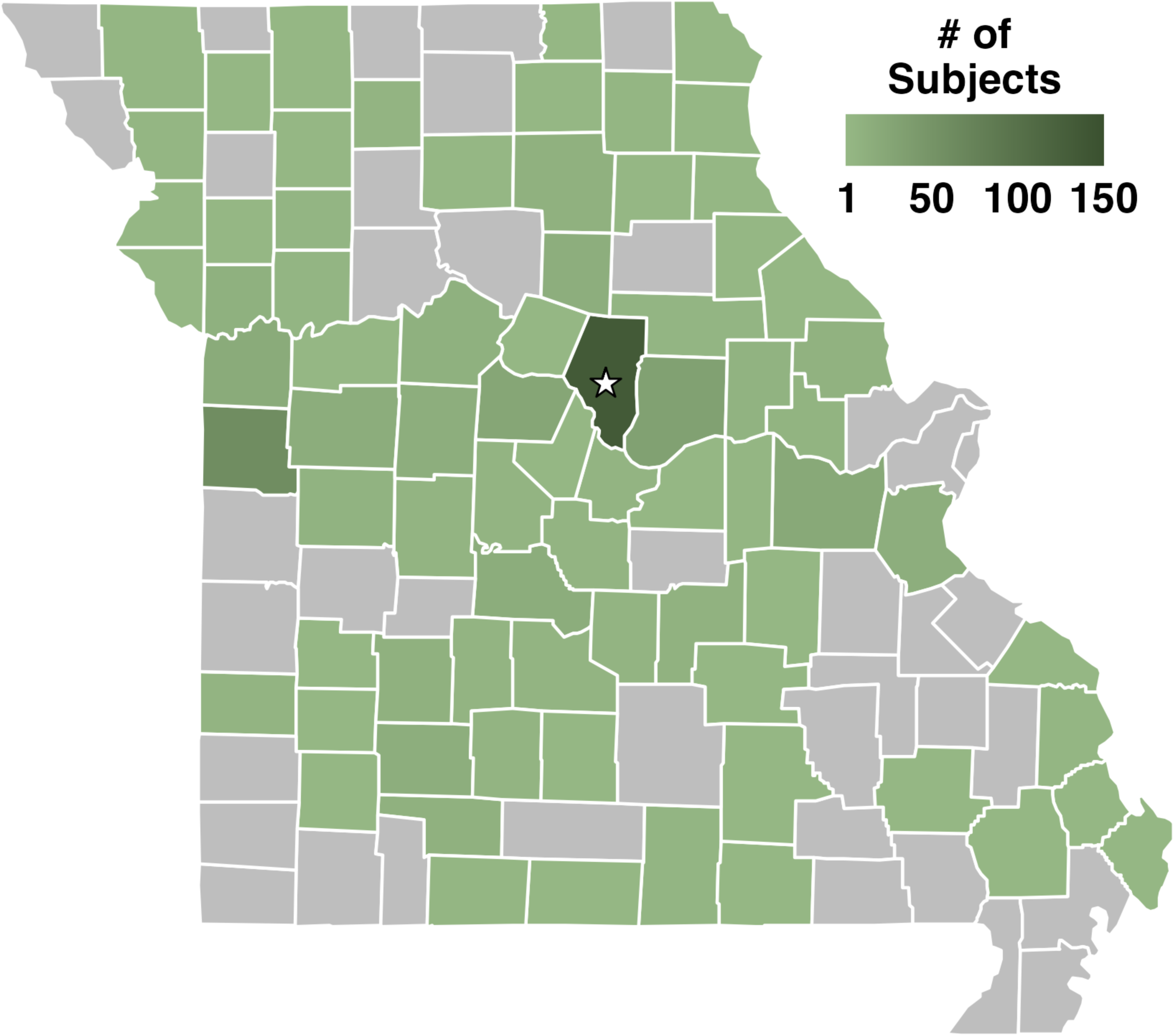
County-level map of the state of Missouri (USA) colored by the number of donating subjects originating from that county. Overall, 68.7% of Missouri counties were represented by at least one patient. The star indicates the University of Missouri which donated a total of 1,218 samples from 707 equids to the EGG database, of which 1,072 samples from 627 equids were collected from Missouri counties.

**Extended Data Fig. 2.**
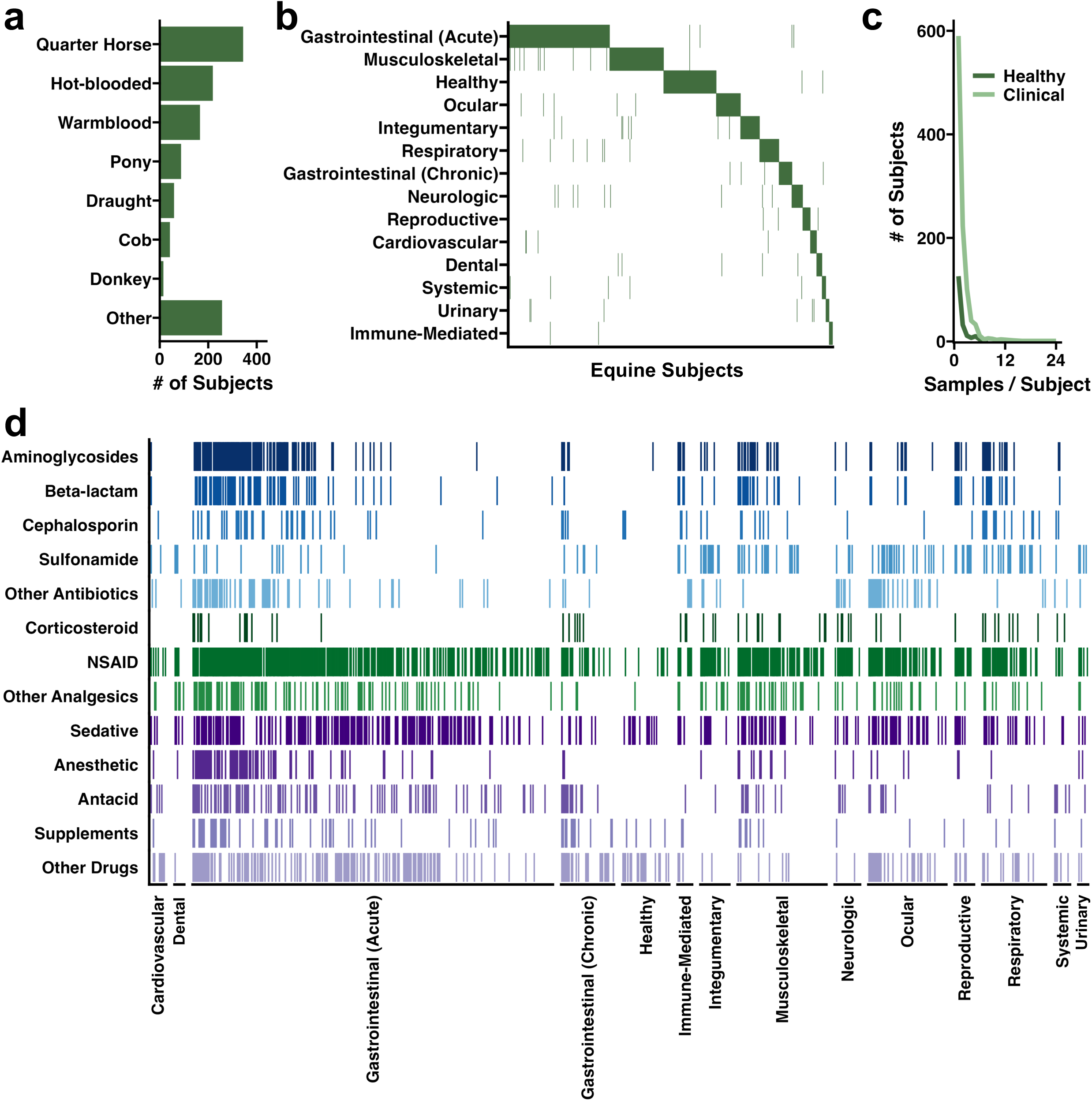
Additional demographic summary of the EGG database. **a**, Bar chart depicting the number of subjects within general breed type. **b**, Tile plot showing all diagnoses recorded for each equine patient. Some subjects came into the hospital for multiple visits. **c**, Line plot depicting the number of subjects with longitudinal data (i.e., multiple samples per visit). **d,** Tile plot depicting classes of drugs (or supplements) given to equine subjects. Columns represent individual subjects.

**Extended Data Fig. 3.**
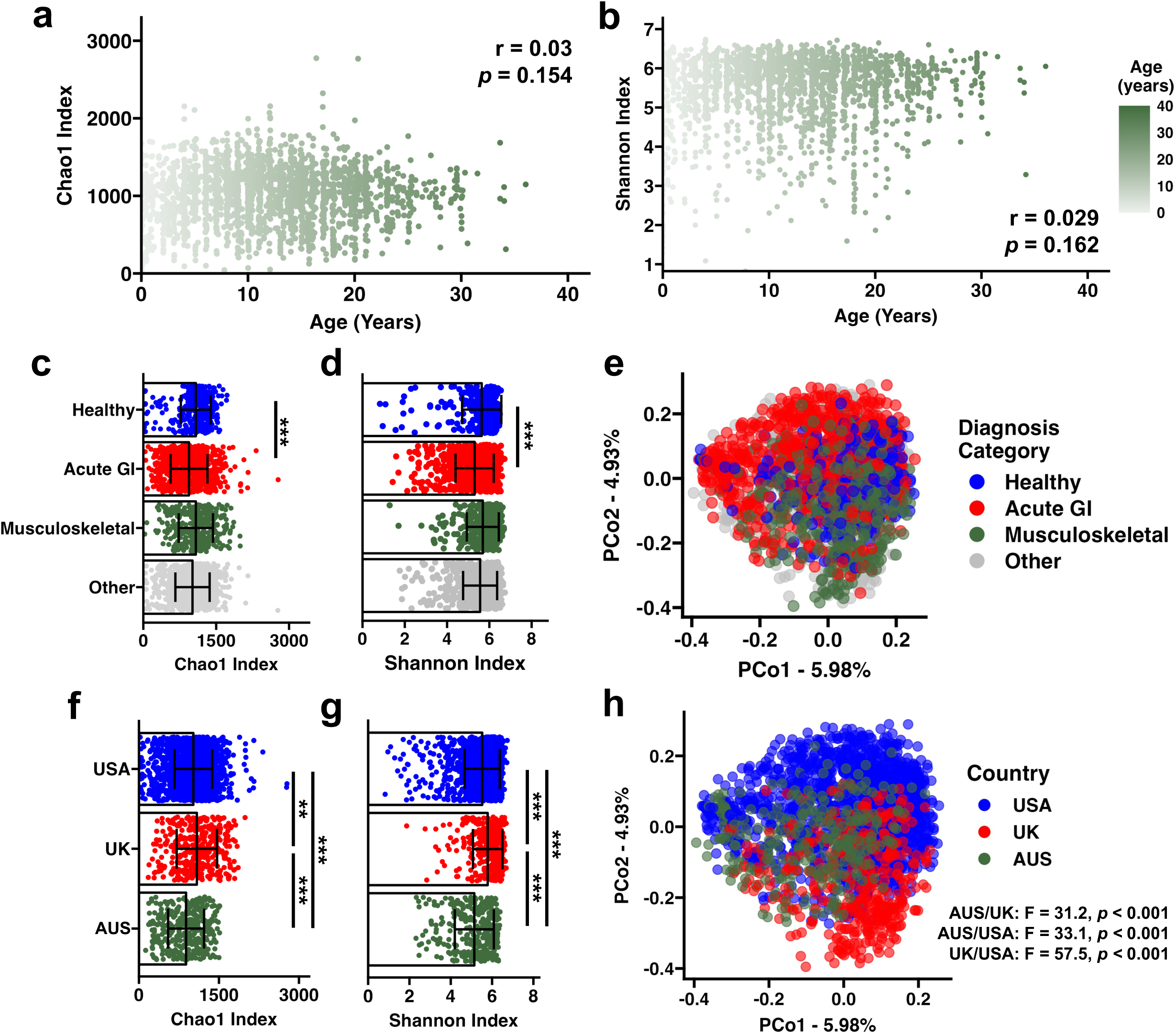
Additional summary of the EGG 16S rRNA dataset. Scatter plots depicting the correlation of sample richness **(a)** or diversity **(b)** and patient age at collection (years). Inset text depicts correlation test using the Spearman method. *N* = 2,302 samples. Dot and bar plots depicting richness **(c)** and diversity **(d)** of the three primary disease categories with the greatest number of samples. All other disease categories were grouped together as “Other.” Bars depict significance between healthy and Gastrointestinal (Acute) groups using pairwise Wilcoxon tests with a Benjamini-Hochberg correction. ** *p* < 0.01, *** *p* < 0.001. All pairwise comparisons, including all fourteen disease categories, may be found in Extended Data Table 2. **e,** PCoA depicting large differences in microbial diversity between healthy, musculoskeletal, and gastrointestinal (acute) samples. All other disease categories were grouped together as “Other.” Statistical comparison between all disease categories may be found in Extended Data Table 2. Dot and bar plots depicting richness **(f)** and diversity **(g)** by geographic location. Bars depict significance between countries using pairwise Wilcoxon tests with a Benjamini-Hochberg correction. ** *p* < 0.01, *** *p* < 0.001. **h,** PCoA depicting large differences in microbial diversity between countries. Inset text depicts pairwise PERMANOVA testing. *N* = 2,362 samples.

**Extended Data Fig. 4.**
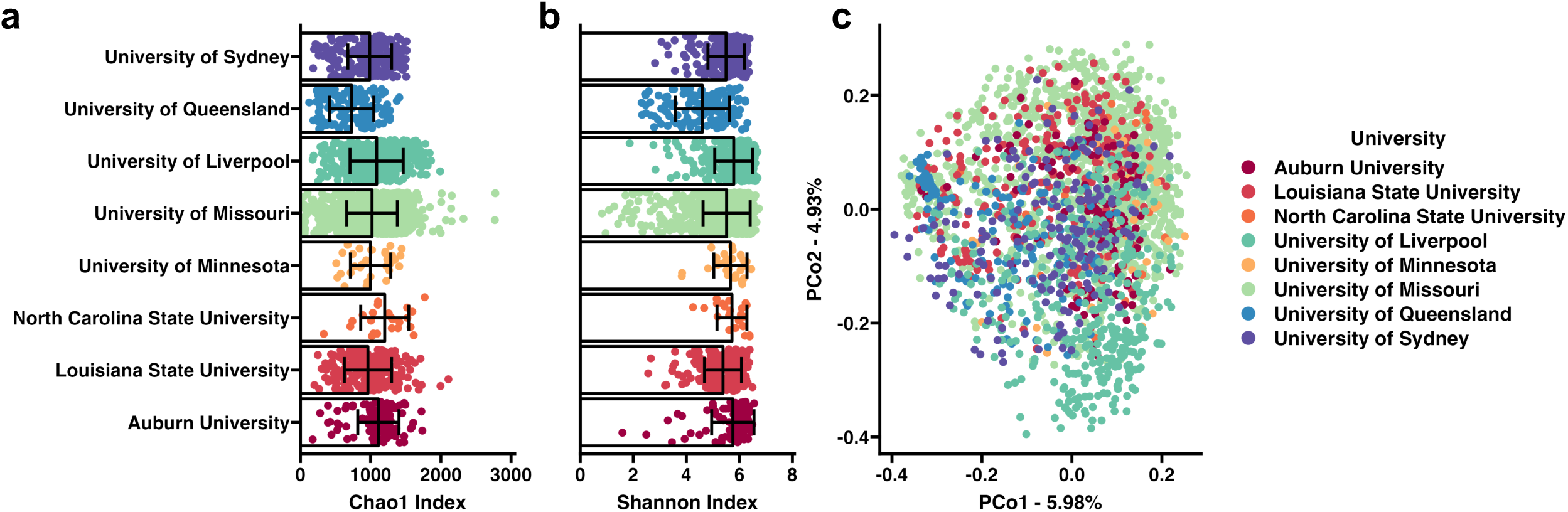
Alpha and beta diversity differences across donating veterinary teaching hospitals. Dot and bar plots depicting the **(a)** richness and **(b)** diversity of samples collected at each donating veterinary teaching hospital. **c,** PCoA depicting differences in beta diversity by donating veterinary teaching hospital. Pairwise comparisons assessing differences in richness, diversity, and microbial composition may be found in **Extended Data Table 3**. *N* = 2,362 samples.

**Extended Data Fig. 5.**
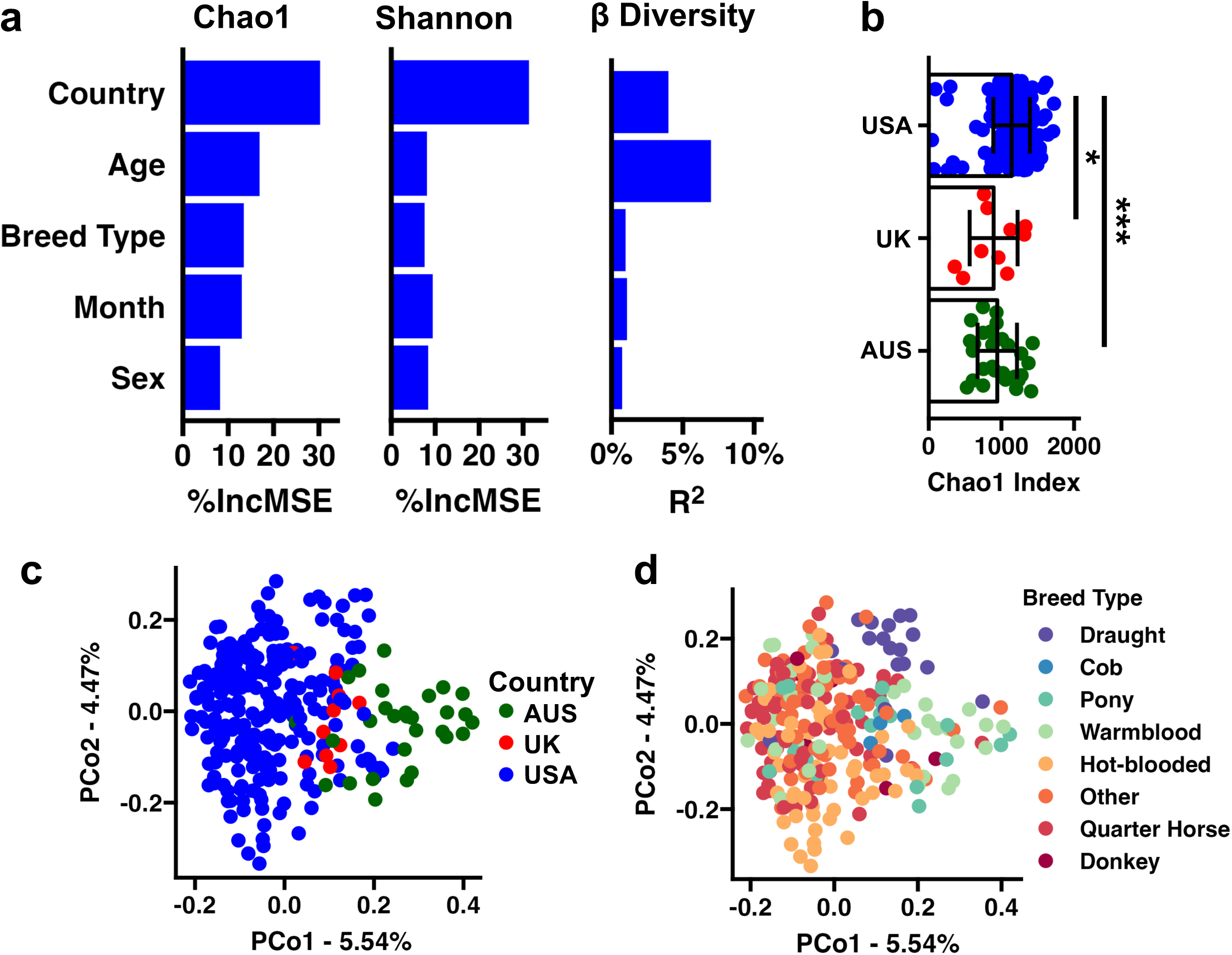
Factors affecting healthy, adult equine microbiome. **a**, Bar plots depicting the importance of various covariates influencing richness *(left)* and diversity *(middle)* and effect size on weighted beta diversity *(right)*. **b**, Dot and bar plot depicting sample richness by country. \**p* < 0.05, \*\*\**p* < 0.001, Wilcoxon Rank-Sum test. PCoA using weighted beta diversity depicting the geographic location **(c)** and breed type **(d)** of healthy subjects.

**Extended Data Fig. 6.**
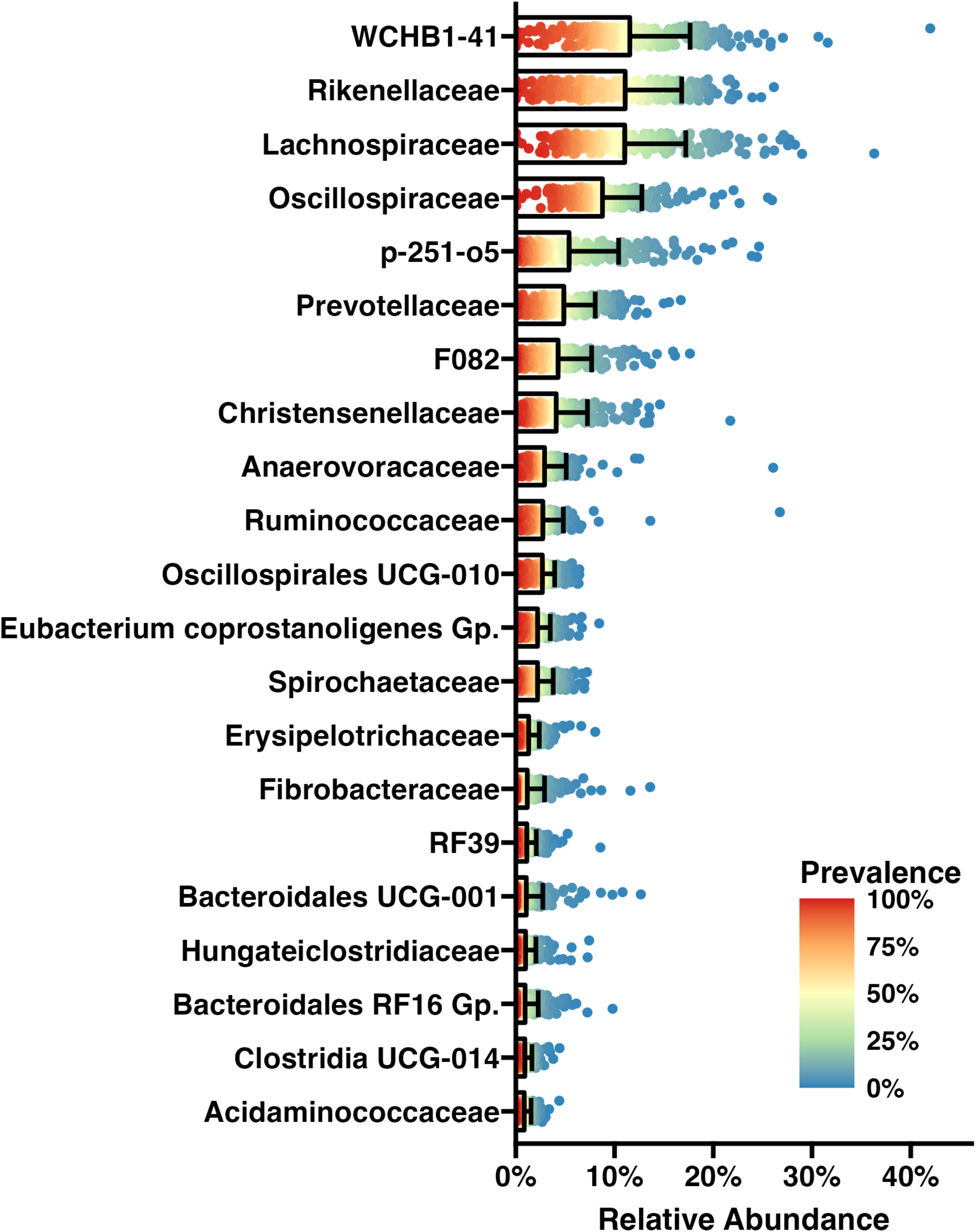
Abundance of core-microbiome. Dot and bar plot depicting the relative abundance of each family detected in the core microbiome of healthy, adult subjects. Color of each point represents the prevalence of that family at that relative abundance threshold (i.e., X% of subjects have at least Y% relative abundance).

**Extended Data Fig. 7.**
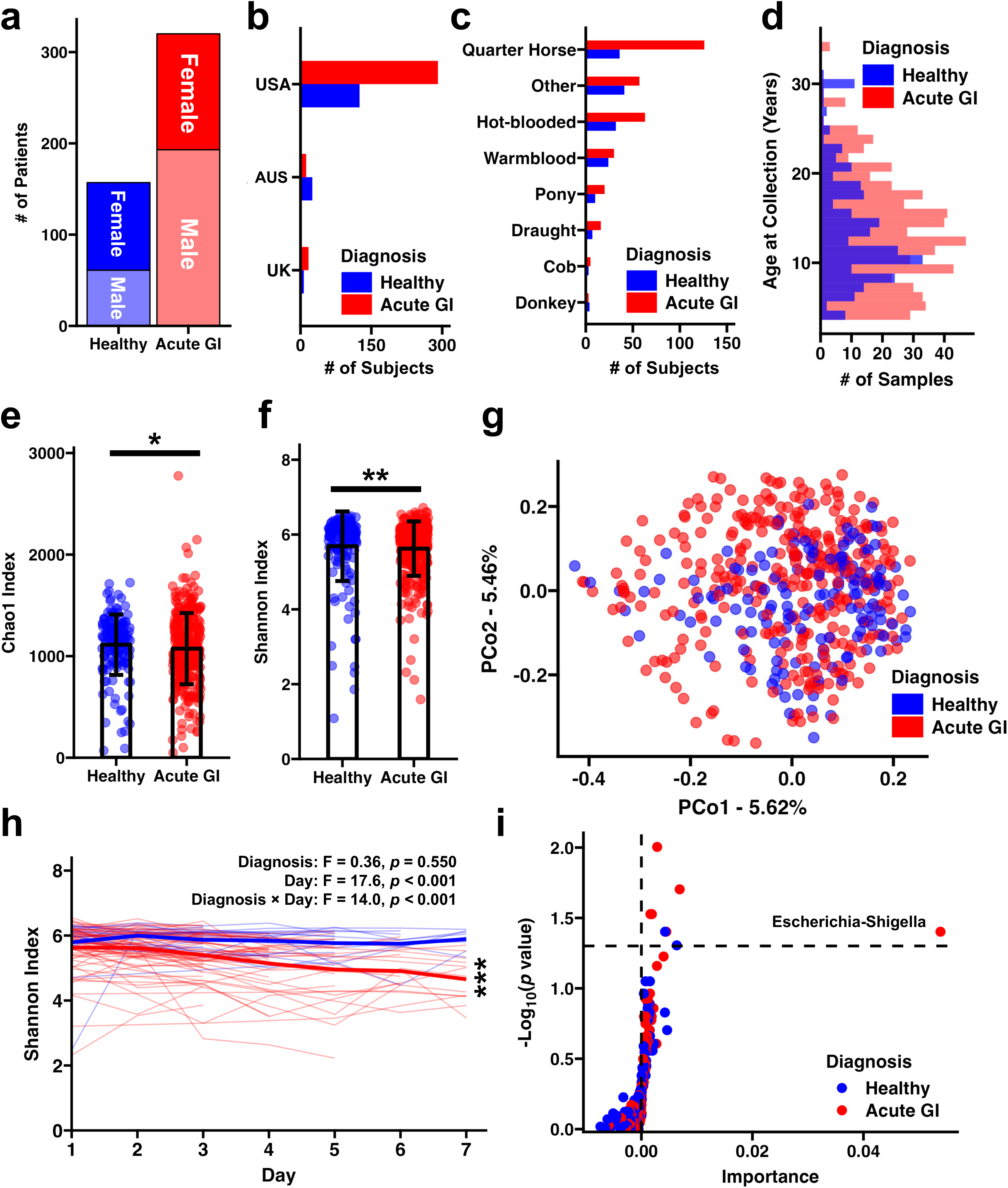
Demographics and microbial differences between healthy and acute GI disease subjects. **a**, Stacked bar plots depicting the number of male and female healthy and acute GI subjects. Bar charts depicting the number of healthy or acute GI disease subjects from each geographic location **(b)** and of each breed type **(c). d,** Histogram depicting the distribution of ages at collection from healthy and acute GI disease populations. Only adults (>4 years old) were used in this analysis. Dot and bar plots showing reduced **(e)** richness and **(f)** diversity in acute GI subjects compared to healthy animals * *p* < 0.05, ** *p* < 0.01. Wilcoxon Rank Sum Test. **g,** PCoA depicting significant differences in weighted beta diversity between healthy and acute GI subjects. F = 3.61, *p* < 0.001. One-way PERMANOVA. **h,** Line plot depicting longitudinal changes in sample diversity (Shannon Index). Thin lines depict individual subjects and thick lines represent group means. Inset text describes two-way ANOVA results of longitudinal data fitted to a LMEM. \*\*\**p* < 0.001. One-way ANOVA within group. **i**, Dot plot depicting the importance and significance of genus-level features in predicting disease status using relative abundance data collected at patient intake.

**Extended Data Fig. 8.**
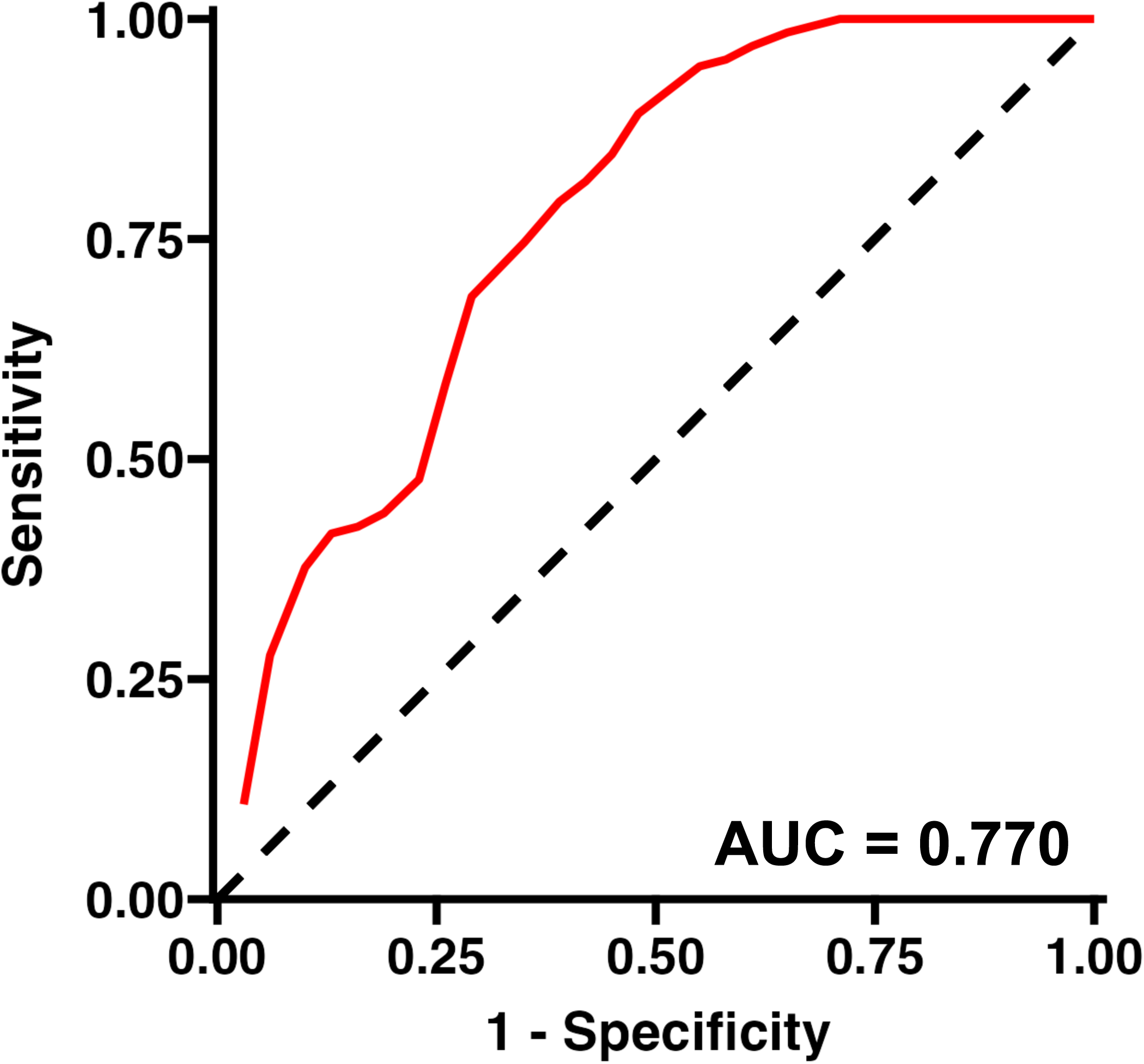
Predicting disease status. Receiver Operating Characteristic (ROC) Curve showing performance of random forest model using genus-level relative abundance data in predicting healthy or acute GI disease status.

**Extended Data Fig. 9.**
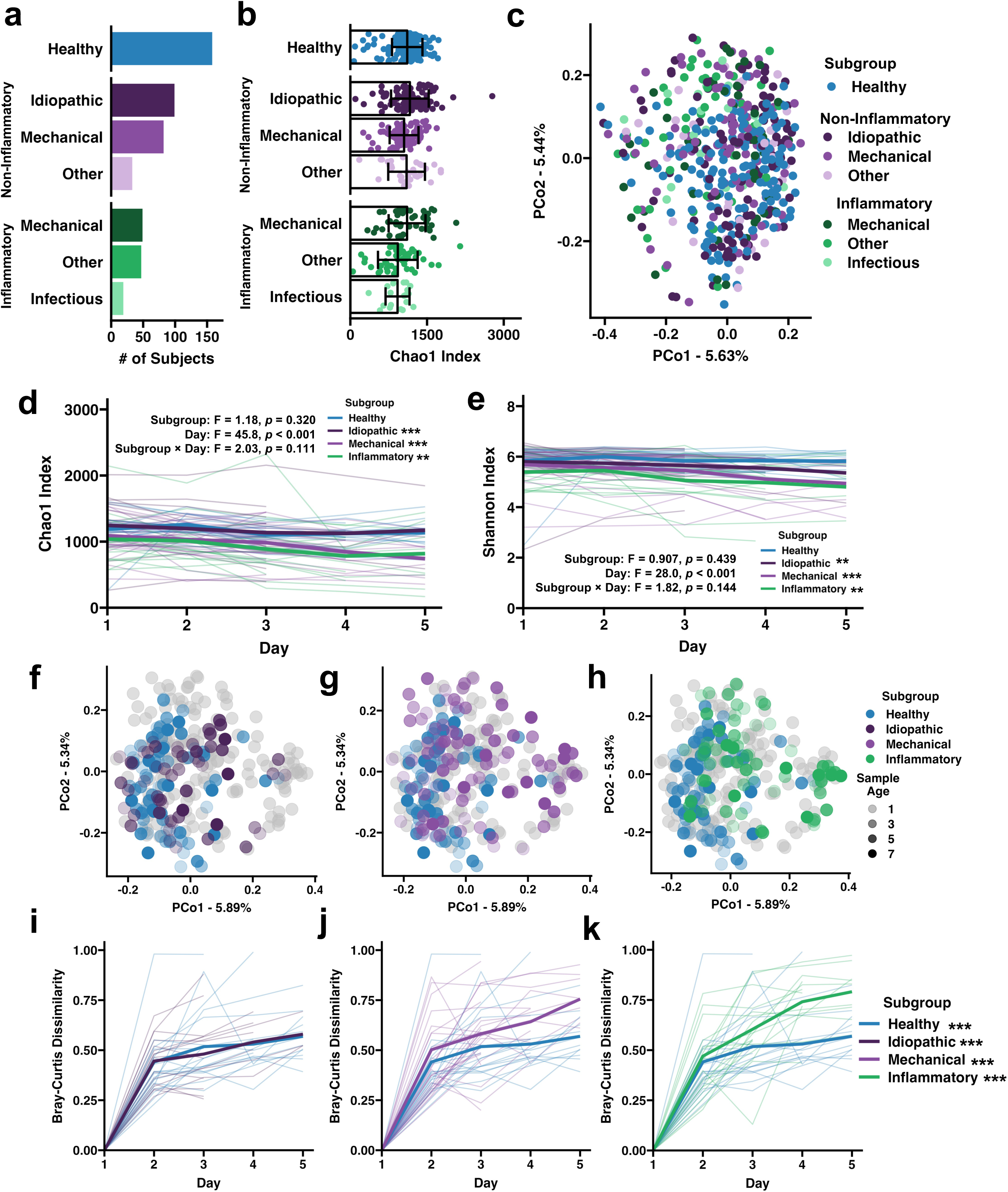
Subgroups of acute GI disease are characterized by microbial dysbiosis. **a,** Bar plot showing the number of subjects per acute GI disease subgroup. **b**, Dot and bar plot depicting sample richness at intake by acute GI disease subgroup. Pairwise comparisons between groups can be found in Extended Data Table 7. **c,** PCoA showing differences in weighted beta diversity between acute GI subgroups at intake. Diagnosis: F = 1.80, *p* < 0.001. One-way PERMANOVA. Pairwise comparisons between groups can be found in Extended Data Table 7. Line plots showing longitudinal changes in sample richness **(d**) and diversity **(e).** Thin lines depict individual subjects, and thick lines represent group means. Inset text describes two-way ANOVA results of longitudinal data fitted to a LMEM. \**p* < 0.05, \*\**p* < 0.01, \*\*\**p* < 0.001. One-way ANOVA within group. **f-h**, PCoA using weighted beta diversity depicting the relationship of healthy and non-inflammatory mechanical **(f)**, inflammatory **(g),** and non-inflammatory idiopathic **(h)** colic subgroups. **i-k**, Volatility assessment of microbial composition comparing weighted dissimilarity of each sample to the sample collected at intake from the same patient between healthy and non-inflammatory mechanical **(i),** inflammatory **(j),** and non-inflammatory idiopathic **(k)** subgroups. Thin lines depict individual subjects, and thick lines represent group means. Inset text describes two-way ANOVA results of longitudinal data (Days 2-7) fitted to a LMEM. \*\*\**p* < 0.001. One-way ANOVA within group.

## References

1. Kauter, A. et al. The gut microbiome of horses: current research on equine enteral microbiota and future perspectives. Anim. Microbiome 1, 14 (2019).

2. Singh, V. et al. Butyrate producers, “The Sentinel of Gut”: Their intestinal significance with and beyond butyrate, and prospective use as microbial therapeutics. Front. Microbiol. 13, 1103836 (2023).

3. O’Reilly, G. C. et al. Characterisation of the Faecal Microbiome of Foals from 0-5 Months of Age and Their Respective Mares across Five Geographic Locations. Front. Biosci.-Elite 14, 22 (2022).

4. Ang, L. et al. Gut Microbiome Characteristics in feral and domesticated horses from different geographic locations. Commun. Biol. 5, 172 (2022).

5. Arnold, C. E. et al. The effects of signalment, diet, geographic location, season, and colitis associated with antimicrobial use or Salmonella infection on the fecal microbiome of horses. J. Vet. Intern. Med. 35, 2437–2448 (2021).

6. Curtis, L., Burford, J. H., England, G. C. W. & Freeman, S. L. Risk factors for acute abdominal pain (colic) in the adult horse: A scoping review of risk factors, and a systematic review of the effect of management-related changes. PLoS ONE 14, e0219307 (2019).

7. Costa, M. C. et al. Comparison of the fecal microbiota of healthy horses and horses with colitis by high throughput sequencing of the V3-V5 region of the 16S rRNA gene. PloS one 7, e41484 (2012).

8. Stewart, H. L. et al. Differences in the equine faecal microbiota between horses presenting to a tertiary referral hospital for colic compared with an elective surgical procedure. Equine veterinary journal 51, 336–342 (2019).

9. Weese, J. S. et al. Changes in the faecal microbiota of mares precede the development of post partum colic. Equine veterinary journal 47, 641–9 (2015).

10. Stewart, H. L. et al. Changes in the faecal bacterial microbiota during hospitalisation of horses with colic and the effect of different causes of colic. Equine veterinary journal 53, 1119– 1131 (2021).

11. Elzinga, S. E., Weese, J. S. & Adams, A. A. Comparison of the fecal microbiota in horses with equine metabolic syndrome and metabolically normal controls fed a similar all-forage diet. J Eq Vet Sci 44, 9–16 (2016).

12. Milinovich, G. J. et al. Microbial ecology of the equine hindgut during oligofructose-induced laminitis. ISME J. 2, 1089–100 (2008).

13. Morrison, P. K. et al. The Equine Gastrointestinal Microbiome: Impacts of Age and Obesity. Frontiers in microbiology 9, 3017 (2018).

14. Coleman, M. C., Whitfield-Cargile, C. M., Madrigal, R. G. & Cohen, N. D. Comparison of the microbiome, metabolome, and lipidome of obese and non-obese horses. PloS one 14, e0215918 (2019).

15. Biddle, A. S., Tomb, J. F. & Fan, Z. Microbiome and Blood Analyte Differences Point to Community and Metabolic Signatures in Lean and Obese Horses. Front. Vet. Sci. 5, 225 (2018).

16. Edwards, J. E. et al. Multi-kingdom characterization of the core equine fecal microbiota based on multiple equine (sub)species. Anim. Microbiome 2, 6 (2020).

17. McKinney, C. A. et al. Assessment of clinical and microbiota responses to fecal microbial transplantation in adult horses with diarrhea. PLoS ONE 16, e0244381 (2021).

18. Topçuoğlu, B. D. et al. mikropml: User-Friendly R Package for Supervised Machine Learning Pipelines. J. open source Softw. 6, 3073 (2021).

19. Donnell, M. M. O. et al. The core faecal bacterial microbiome of Irish Thoroughbred racehorses. Lett. Appl. Microbiol. 57, 492–501 (2013).

20. Costa, M. C. et al. Characterization and comparison of the bacterial microbiota in different gastrointestinal tract compartments in horses. Vet. J. 205, 74–80 (2015).

21. Huttenhower, C. et al. Structure, Function and Diversity of the Healthy Human Microbiome. Nature 486, 207–214 (2012).

22. Bulmer, L. S. et al. High-starch diets alter equine faecal microbiota and increase behavioural reactivity. Sci. Rep. 9, 18621 (2019).

23. Baraille, M. et al. Changes of faecal bacterial communities and microbial fibrolytic activity in horses aged from 6 to 30 years old. PLOS ONE 19, e0303029 (2024).

24. Lindenberg, F. et al. Development of the equine gut microbiota. Sci. Rep. 9, 14427 (2019).

25. Mady, E. A. et al. Relationship between the components of mare breast milk and foal gut microbiome: shaping gut microbiome development after birth. Vet. Q. 44, 1–9 (2024).

