## Supplementary material for "A novel dataset of 2,362 equine fecal microbiomes from eight veterinary teaching hospital on three continents reveals dominant effects of geography, breed, and disease": Methods

**Extended Data Methods**

*Participating Institutions*

Samples were collected by veterinarians or staff working in the equine hospitals at the University of Missouri (MU), Auburn University, Louisiana State University (LSU), North Carolina State University (NCSU), and the University of Minnesota (UMN) in the United States (US); the University of Liverpool in the United Kingdom (UK); and the University of Queensland and the University of Sydney in Australia (AUS). Eighty-three samples were collected from a private practice in Cass County, Missouri, US in collaboration with the MU Veterinary Teaching Hospital.

*Patients*

Patients included in the current study were admitted to one of eight participating institutions for reasons related to new or ongoing health concerns, or for reasons unrelated to adverse health including reproductive exams, prophylactic dental care, or accompanying a foal. Details regarding patient demographics, health status, treatment, and outcomes were manually gathered by each contributing institution and sent to the MU for curation in a uniform format. Animals older than four years of age were considered adults. The following terms were used to classify male and female equids: fillies (female, < 4 years old), colts (male, < 4 years old), geldings (castrated adult males), stallions (intact adult males), mares (adult females), jacks (adult male donkeys), and jennies (adult female donkeys).

*Sample Collection, Storage, and Transport*

Freshly evacuated fecal samples were collected from the stall floor, following removal of visible bedding material, dust, or other material from the exterior of the fecal bolus. Following admission to a participating equine hospital, sample collection was attempted daily until case resolution or discharge from the hospital. Typically, samples were collected within a one- to two-hour window during the morning, immediately after stalls had been cleaned and replaced with clean bedding. Fecal boli present in freshly changed bedding were collected into sterile specimen containers, labeled with the patient ID and date, and placed in a -20°C or -80°C freezer located in the hospital. MU samples were then periodically transported in insulated containers containing freezer gel-packs, by car, to the MU Metagenomics Center for DNA extraction.

At other participating institutions, fecal samples were collected into 50 mL conical tubes (ThermoFisher Scientific) and kept frozen until shipping directly to the MU Metagenomics Center for DNA extraction. Samples were shipped *en masse* in insulated shipping containers containing dry ice via World Courier for shipments from LSU, Auburn, NCSU, UMN, and UK, and FedEx for shipments from AUS institutions. All international shipments were approved by US Customs and accompanied by full declarations. Couriers periodically opened packages to replenish dry ice during transport of international shipments, and continuous temperature monitors packed with those samples confirmed that all shipments remained frozen throughout transport.

*DNA Extraction*

All DNA extractions were performed at the MU Metagenomics Center (Columbia, MO, US). DNA was extracted from fecal samples (≤ 250mg) in groups of 24 samples, using QIAamp PowerFecal Pro DNA Extraction Kits (Qiagen), per the manufacturer’s instructions with the following exception. During the initial bead-beating process, tubes were agitated using a Qiagen TissueLyser II at maximum speed (30 Hz) for ten minutes, rather than the vortex adapter described in the manufacturer’s protocol. Following extraction, DNA yields were determined using Quant-iT dsDNA Assay Kits, broad range (BR) with a Qubit 2.0 fluorometer (Invitrogen).

*16S rRNA Amplicon Library Preparation and Sequencing*

Library preparation and sequencing were performed at the MU Genomics Technology Core. Bacterial 16S rRNA amplicons were constructed via amplification of the V4 region of the 16S rRNA gene with universal primers (U515F/806R), flanked by Illumina standard adapter sequences^1,2^. Dual-indexed forward and reverse primers were used in all reactions. PCR was performed in 50 µL reactions containing 100 ng metagenomic DNA, primers (0.2 µM each), dNTPs (200 µM each), and Phusion high-fidelity DNA polymerase (1U, Thermo Fisher). Amplification parameters were 98°C^(3 min)^ + [98°C^(15 sec)^ + 50°C^(30 sec)^ + 72°C^(30 sec)^] × 25 cycles + 72°C^(7 min)^. Amplicon pools (5 µL/reaction) were combined, thoroughly mixed, and then purified by addition of Axygen Axyprep MagPCR clean-up beads to an equal volume of 50 µL of amplicons and incubated for 15 minutes at room temperature. Products were then washed multiple times with 80% ethanol and the dried pellet was resuspended in 32.5 µL EB buffer (Qiagen), incubated for two minutes at room temperature, and then placed on the magnetic stand for five minutes. The final amplicon pool was evaluated using the Advanced Analytical Fragment Analyzer automated electrophoresis system, quantified using quant-iT HS dsDNA reagent kits, and diluted according to Illumina’s standard protocol for sequencing as 2×250 bp paired-end reads on an Illumina MiSeq instrument.

*Bioinformatics*

16S rRNA sequences were processed using the Quantitative Insights into Molecular Ecology 2 (QIIME2) v2024.5^3^ framework in multiple batches. Paired-end reads were trimmed of the universal primers and Illumina adapters using *cutadapt*^4^. Samples yielding less than 10,000 forward reads were discarded. Reads were then denoised into unique amplicon sequence variants (ASVs) using DADA2^5^ with the following parameters: 1) reads were truncated to 150 bp in length, 2) reads with greater than 2 expected errors were discarded, 3) reads were merged with minimum overlap of 12 bp, 4) chimeras were removed using the ‘consensus’ method. Unique sequences were filtered to between 249 and 257 bp in length. Unique sequences were then assigned taxonomy using an a *sklearn* algorithm and a 99% non-redundant SILVA v138 reference database^6^ trimmed to the 515F/806R^2^ region of the 16S rRNA gene. As batches of samples were sequenced, filtered feature tables, representative sequences, and taxonomic classifications were merged within QIIME2.

*Statistical Analysis*

All data analyses and visualizations were performed using available libraries within R^7^ v4.2.2. A *p* value of less that 0.05 was considered significant. Visualizations were created using various tools within the *ggplot2* library^8^. All code may be accessed at <https://github.com/ericsson-lab/EquineGutGroup>.

Univariate data including alpha diversity and dissimilarity values were tested for normality using the Shapiro-Wilkes test from the *rstatix* library^9^. For single timepoint analyses, differences in non-normally distributed data were determined using the Kruskal-Wallis test from the *rstatix* library^9^. *Post hoc* testing was performed using pairwise Wilcoxon Rank Sum tests with Benjamini-Hochberg corrections^9,10^ for multiple comparisons, when appropriate. Longitudinal comparisons were made by fitting longitudinal data to linear mixed effects (LME) models using the *lme4*^11^ and *lmerTest*^12^ library accounting for multiple measurements of individual animals: *outcome ~ covariates_of_interest + (1 | patient_id)*. The effects of time and grouping variable of interest were then assessed using one- or two-way analysis of variance (ANOVA) tests. Estimated means of linear trends analyses from the *emmeans* library^13^ were used to estimate the slope of changes over time within groups using the following models: *LME_model* *~ grouping_variable | sample_age*.

Differences in multivariate data were assessed using permutational multivariate analysis of variance (PERMANOVA) testing with weighted (Bray-Curtis) distances. Using the *adonis2* function within the *vegan* library^14,15^, one- and two-way PERMANOVA testing was performed using 9,999 permutations. Effect size (R^2^) of selected covariates on beta diversity were extracted from PERMANOVA testing. Pairwise one-way PERMANOVA testing was performed using the *pariwise.adonis2* function from the *pairwiseAdonis*^16^ library with 9,999 permutations. Differences in beta diversity were visualized using principal coordinate analyses (PCoA). PCoA were performed using the *ape*^17^ library with a Cailliez correction. The first two principal coordinates of each PCoA were visualized using *ggplot2*^8^*.*

The core microbiome analysis was performed using the *plot_core* function from the *phyloseq* library^18^. The parameters for the core microbiome analysis were defined as families detected with a prevalence in 5% of samples at a minimum relative abundance 1%. The prevalence of taxa was determined up to a minimum of 10% relative abundance. Relative abundance data for each feature of the core microbiome was then fitted to LME models and the effects of select covariates were determined using ANOVA testing. Effect sizes (partial eta squared) were extracted using the *eta_squared* function from the *effectsize* library^19^.

Host and environmental factors influencing richness and diversity measurements were determined using Breiman's random forest regression algorithm from the *randomForest* library^20^. All samples with complete metadata were included in the training set to determine importance of general metadata factors on richness and diversity. Models were trained using 1,000 trees. Importance values (i.e., % increase in mean squared error) were extracted using the *importance* function.

A random forest classifier predicting health status (healthy or acute gastrointestinal disease) was created using the *mikropml* library^21^. Genus-level relative abundance data was preprocessed removing uninformative features with near-zero variance and transforming data using the default “center” and “scale” methods. The *run_ml* function from the *mikropml* library^21^ was then used to generate a random forest classifier to predict health status using the preprocessed genus-level relative abundance data. Default parameters including five-fold cross-validation with 100 partitions were used to build the model. Data were split 80/20 for training and testing, respectively. Feature importance was determined by taking the difference of the true model performance (i.e., area under the curve [AUC] of model receiver operating characteristic [ROC] curve) and model performance when that feature was removed. Significance was determined using permutation testing of feature importance. The random seed was set to 1851 for reproducibility.

**References**

1. Walters, W. A. *et al.* PrimerProspector: de novo design and taxonomic analysis of barcoded polymerase chain reaction primers. *Bioinformatics* **27**, 1159–61 (2011).

2. Caporaso, J. G. *et al.* Global patterns of 16S rRNA diversity at a depth of millions of sequences per sample. *Proc. Natl. Acad. Sci.* **108**, 4516–4522 (2011).

3. Bolyen, E. *et al.* Reproducible, interactive, scalable and extensible microbiome data science using QIIME 2. *Nat Biotechnol* **37**, 852–857 (2019).

4. Martin, M. Cutadapt removes adapter sequences from high-throughput sequencing reads. *EMBnetJ.* **17**, 10–12 (2011).

5. Callahan, B. J. *et al.* DADA2: High-resolution sample inference from Illumina amplicon data. *Nature methods* **13**, 581–3 (2016).

6. Quast, C. *et al.* The SILVA ribosomal RNA gene database project: improved data processing and web-based tools. *Nucleic Acids Res.* **41**, D590-6 (2013).

7. Team, R. D. C. R: A Language and Environment for Statistical Computing. (2022).

8. Wickham, H. ggplot2, Elegant Graphics for Data Analysis. 11–31 (2016) doi:10.1007/978-3-319-24277-4_2.

9. Kassambara, A. rstatix: Pipe-Friendly Framework for Basic Statistical Tests. (2022).

10. Benjamini, Y. & Hochberg, Y. Controlling the False Discovery Rate: A Practical and Powerful Approach to Multiple Testing. *J Royal Statistical Soc Ser B Methodol* **57**, 289–300 (1995).

11. Bates, D., Mächler, M., Bolker, B. & Walker, S. Fitting Linear Mixed-Effects Models Using lme4. *J. Stat. Softw.* **67**, (2015).

12. Kuznetsova, A., Brockhoff, P. B. & Christensen, R. H. B. lmerTest Package: Tests in Linear Mixed Effects Models. *J. Stat. Softw.* **82**, (2017).

13. Lenth, R. emmeans: Estimated Marginal Means, aka Least-Squares Means. <https://CRAN.R-project.org/package=emmeans> (2023).

14. Dixon, P. VEGAN, a package of R functions for community ecology. *J Veg Sci* **14**, 927–930 (2003).

15. Oksanen, J. *et al.* vegan: Community Ecology Package. (2022).

16. Arbizu, M. & P. pairwiseAdonis: Pairwise multilevel comparison using adonis. <https://github.com/pmartinezarbizu/pairwiseAdonis> (2020).

17. Paradis, E. & Schliep, K. ape 5.0: an environment for modern phylogenetics and evolutionary analyses in R. *Bioinformatics* **35**, 526–528 (2019).

18. McMurdie, P. J. & Holmes, S. phyloseq: An R Package for Reproducible Interactive Analysis and Graphics of Microbiome Census Data. *PLoS ONE* **8**, e61217 (2013).

19. Ben-Shachar, M., Lüdecke, D. & Makowski, D. effectsize: Estimation of Effect Size Indices and Standardized Parameters. *J. Open Source Softw.* **5**, 2815 (2020).

20. Liaw, A. & Wiener, M. Classification and Regression by randomForest. *R News* (2002).

21. Topçuoğlu, B. D. *et al.* mikropml: User-Friendly R Package for Supervised Machine Learning Pipelines. *J. open source Softw.* **6**, 3073 (2021).
